# Variability in Fruit Yield and Quality of Genetically Diverse Tomato Cultivars in Response to Different Biochars

**DOI:** 10.1101/2020.06.28.176487

**Authors:** Elvir Tenic, Daylen Isaac, Rishikesh Ghogare, Amit Dhingra

## Abstract

**Background:** Intensive agricultural practices have reduced soil health thereby negatively impacting crop yields. There is a need to maintain healthy soils and restore marginal lands to ensure efficient food production. Biochar, a porous carbon-rich material generated from pyrolysis of various feedstock sources is receiving attention as a soil amendment that has the potential to restore soil organic carbon content and also enhance crop yields. However, the physical and chemical properties of biochar are influenced by pyrolysis parameters. These in turn determine its interaction with the soil, thereby influencing its biological properties in terms of impact on soil microcosm and plant productivity. While most studies report the evaluation of one biochar and a single plant cultivar, the role of the plant’s genetic background in responding to biochar as a soil amendment remains unanswered. The impact of six distinct biochars on agronomic performance and fruit quality of three genetically diverse tomato (*Solanum lycopersicum*) cultivars was evaluated to test the hypotheses that 1) biochars derived from different feedstock sources would produce unique phenotypes in a single cultivar of tomato, and 2) single feedstock-derived BC would produce different phenotypes in each of the three tomato cultivars.

**Results:** Different biochars impacted shoot dry weight, total fruit weight, and yield per plant in each cultivar differently. Both positive and negative effects were observed depending on the biochar-cultivar combination. In ‘Oregon Spring’, Ryegrass straw and CoolTerra biochar enhanced yield. In ‘Heinz’, an increase in fruit weight and citric acid was observed with several of the biochars. In ‘Cobra’, improved yields were accompanied by reduction in fruit quality parameters. Both hypotheses were supported by the data.

**Conclusions:** This study demonstrated that the genetic background of a plant is an important variable in determining the outcome of using biochar as a soil amendment. Strategies for application of biochar in agricultural production should consider the variables of soil type, feedstock source, pyrolysis parameters and plant genetic background for enhancing crop productivity and carbon sequestration.

## 1. Background

Agricultural soils have been strained to reach their highest potential in productivity and now encounter several biotic and abiotic challenges. Years of intensified crop production has adversely impacted soil health. Ever-increasing application of fertilizer and irrigation has resulted in the loss of organic matter and sodification, leading to deterioration of soil tilth [1]. To combat these impending detriments to soil health, management approaches are being adopted to increase soil organic matter (SOM), foster a diverse soil microcosm, improve crop productivity, and promote additional ecosystems services [2–5]. However, due to changing climatic conditions, soil organic carbon (SOC) levels are projected to decrease in the future [6]. Therefore, it is critical to pursue interventions that encourage beneficial soil practices such as implementing cover crops and reduced tillage [7–9]. Such measures will aid in the development of carbon negative ecosystems, which focus on returning carbon assimilated by plants back into the soil in a stable form with a long half-life. The carbon positive cycle promoted by intense agriculture has further heightened the challenges posed by soil erosion and changing climate conditions [10,11]. These challenges need to be addressed to secure global food supplies for the current and future generations.

The practices of early indigenous cultures of the Amazon over 2500 years ago are known to have improved soil health through incorporation of burnt biomass, resulting in production of a high-fertility ‘*Terra Preta’* layer atop the otherwise sub-fertile Amazon soil [3,12–15]. In the 17^th^ century, Japanese agriculturalists experimented with low-oxygen burnt rice husk as a soil amendment for more intense crop production [16]. Recently, there has been an emerging emphasis on the use of burnt biomass, or biochar, to reclaim the health of marginal soils.

Biochar (BC) is a carbon-rich, porous product generated by a thermochemical process known as pyrolysis. It involves controlled burning of feedstock under low oxygen levels at temperatures ranging from 300°C to 800°C [17,18]. Production of biochar can be achieved using various feedstocks, the most common of which include agricultural crop residue, organic manure, and wood [19]. With improvements in automation, and with growing knowledge of the utility of BC as a soil amendment with the potential to enhance nutrient availability and facilitate long-term carbon sequestration, it is now feasible to produce consistent quality biochar that is expected to spur its utilization both in research and farming [20–24].

Specific impacts of biochar amendment to soil include alterations in bulk density, porosity, and water retention; these properties make the exchange of water, nutrients, and gases more efficient, resulting in enhanced crop productivity [25,26]. Additionally, since BC is a stable source of carbon and nutrients, it influences the soil microcosm by fostering the proliferation of microbial communities for extended periods, which in turn enhance soil tilth and health [27]. The biological, chemical, and physical influence of biochar and its role in enhancing soil health is well-documented; however, its utilization in soils produces a spectrum of outcomes in terms of crop productivity [28–34].

Productivity in a diverse range of crops, like tomato, lettuce and other leafy vegetables, beans, potato, wheat, maize, and rice to name a few, has been evaluated in soils amended with biochar derived from various feedstocks [30,35–40]. The feedstock source determines the final nutrient profile of the biochar. Organic waste feedstocks generate biochar rich in potassium and phosphorus, low in C levels, and low in surface area. Biochar derived from wood feedstocks is enriched in organic matter and surface area; however, it has low N, P, and K levels, and reduced capacity for cation exchange. Generally, crop residue-derived biochars are rich in N [41–43]. The variation in nutrient profiles along with other physical properties determines how the biochar interacts with the soil and collectively influences plant performance.

Several recent meta-analyses of the various studies investigating the role of biochar on crop productivity conclude that, overall, there is a positive impact on crop yield [22,43,44]. However, there are studies where biochar amendment impacts one aspect of plant development but has no impact on yield or it produces a detrimental outcome [32,37,45]. It is well-known that the genetic background of a plant influences how it responds to a given stimulus [46–48]. Interestingly, most previous reports evaluating the impact of biochar have studied one cultivar’s response to biochar derived from a single feedstock. The question then emerges of whether different cultivars will respond differently to biochars derived from different feedstocks.

In this study, the impact of biochars derived from six different feedstocks on the growth and development of three genotypically-distinct cultivars of tomato (*Solanum lycopersicum* L.) was evaluated. Experiments were conducted to test the following hypotheses: 1) Biochars derived from different feedstock sources will produce unique phenotypes in a single cultivar of tomato, and 2) a single feedstock-derived biochar will produce different phenotypes in each of the three tomato cultivars.

## 2. Methods

### a. Biochar source

Five types of BC generated from their respective feedstocks were provided by Ag Energy Systems, (Spokane, WA). The feedstocks used were as follows: Ryegrass straw (RGS), Ryegrass tailings (RGT), Russian thistle (RT), thermomechanical pulp waste (TMP), and Walnut shell (W). A commercially available BC product, Cool Terra® (CT), manufactured by Cool Planet (Greenwood Village, CO), was also used in the study. All experiments were conducted with 0.5% and 1% w/w rates of BC amendment.

### b. SEM and EDX Analysis

Scanning Electron Microscopy (SEM) was performed on each biochar at the Franceschi Microscopy and Imaging Center at Washington State University. A sample of each biochar was fixed to a pin stub and sputter coated in gold. SEM samples were imaged on a Tescan Vega SEM equipped with an Energy-dispersive X-ray spectroscopy (EDX) detector in order to make a qualitative visual assessment of porosity and general particle size. Qualitative elemental composition data for each biochar was collected with the EDX detector.

### c. Plant growth conditions

Three cultivars of tomato (*Solanum lycopersicum* L.) representing unique market applications and diverse genetic backgrounds were selected for this experiment. ‘Oregon Spring’, an heirloom determinate variety was selected due to its popularity in home gardening. ‘Heinz 2653’, also a determinate variety, is commonly used as a commercial processing tomato. ‘Cobra F1’, an indeterminate variety, was selected due to its commercial use as a greenhouse variety.

Seeds for the three cultivars were obtained from Territorial Seed Company (Cottage Grove, Oregon). Seeds were germinated in 4-inch rockwool squares and grown to 4-5 nodes (15-20 cm) in height. Afterwards, plantlets were transplanted into 2.8 L pots with either organic Sunshine Mix#1/LC1 (Sun Gro Horticulture, Massachusetts) as a control or Sunshine Mix containing biochar (BC) at 0.5% or 1% (w/w) rate. One week after transplant, each pot was fertilized twice a week with 450 mL of dilute (20 mL/L water) organic Alaska 5-1-1 Fish Fertilizer (Lilly Miller Brands, CA). Plants were maintained in the glasshouse at the Washington State University Plant Growth Facilities with temperatures held at 24°C/18°C (day/night); relative humidity was maintained at 40-60%. High Pressure Sodium (HPS) lights provided supplemental lighting, extending day length to 16 hours as needed. Young plants were watered every other day, while the larger, mature plants were watered daily.

### d. Experimental Design

Six individual experiments were conducted, with two experiments each for Oregon Spring’, ‘Heinz 2653’ (‘Heinz’), ‘and ‘Cobra’ F1, as reported in Table 1A and B. Experiments with ‘Heinz’ and ‘Oregon Spring’ were conducted over a period of 102 days while with ‘Cobra’ F1 over a period of 182 days. All six experiments were conducted independently with randomized design in the Washington State University Glasshouse. Each experiment consisted of 56 plants: eight plants contained 0% BC and served as controls, while four plants were randomly assigned to each of the 13 treatment groups (Table 1B).

**Table 1A.** Planting and harvest dates for each experiment with three tomato cultivars.

| Experiment | Cultivar | Date Planted | Date Harvested |
| --- | --- | --- | --- |
| 1 | Oregon Spring | 2/17/17 | 6/3/17 |
| 2 | Oregon Spring | 1/20/18 | 5/7/18 |
| 1 | Heinz | 2/17/17 | 6/3/17 |
| 2 | Heinz | 5/15/17 | 8/30/17 |
| 1 | Cobra | 5/16/17 | 11/10/17 |
| 2 | Cobra | 11/8/17 | 5/9/18 |

**Table 1B.**
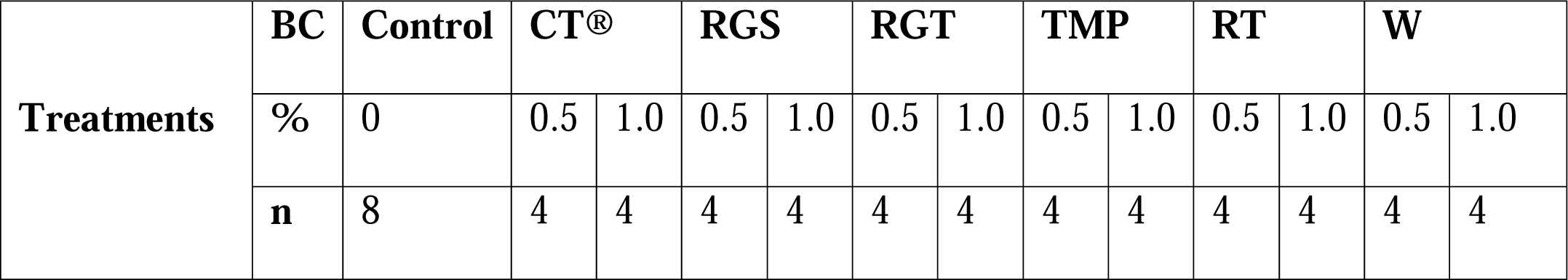
Experimental design for biochar treatments and number of plants used for each. CT – Cool Terra®, RGS – Ryegrass Straw, RGT – Ryegrass tailings, TMP – Thermomechanical pulp, RT – Russian thistle, W – Walnut.

### e. Plant growth parameters and assessment of fruit quality

Dry Weight: Aboveground shoot biomass was collected at the conclusion of each experiment. Fruits were removed, plants were cut at soil level to remove the roots, and the shoots were completely dried in large paper bags at 60°C for 48 hours prior to weighing.

Yield: To measure yield, four random fruits per plant were selected for sampling at ‘Breaker’ stage, which is defined as the point in developmental where less than 10% of surface area displays color change [49]. Following achievement of the ‘Red’ stage, the point in development where greater than 90% of a fruit’s surface area displays color change, fruit were collected at regular intervals throughout the remainder of the experiment [49]. The yield for each plant was quantified based on the total number of fruits and cumulative fruit weight.

Quality: Fruit quality was assessed by quantifying total soluble solids, sugars, and organic acid content. Briefly, a handheld rotary Bio-Homogenizer (model M133/1281-0 from Biospec Products Inc. Bartlesville, OK) was used to extract juice from five grams of fruit (flesh and peel tissue) from each of the four sampled fruit. Juice extracted from ‘Red’ stage fruit was filtered through cheese cloth and used for refractometer-based quantification of total soluble solids (TSS). An aliquot of the juice sample was quickly centrifuged, the resulting supernatant was filtered using 0.45 µm pore size filters, and the sample stored at −80°C for later use in quantification of sugar and organic acid profiles. Fructose, glucose, citric acid, malic acid, and fumaric acid were quantified using a Varian Prostar 230 HPLC equipped with an Aminex HPX 87H column coupled to a refractive index (RI) and UV (210 nm) detector. The column was eluted with 0.005M H_2_SO_4_ at a flow rate of 0.6 mL/min at 65°C [50]. Identification and quantification of sugars and organic acids was done by the external standard method [50].

### f. Statistical analysis

All experiments were assessed independently. Data was analyzed with Rstudio (Version 1.1.463) utilizing the Ggplot2 (Version 3.3.0), Tidyverse (Version 1.2.1), and Ggpubr (Version 0.2.3) packages. Significance was tested using pairwise t-tests with α set for 0.1, .05, and 0.01. Figures with one, two, and three stars represent significance at p-value of <0.1, <0.05 and <0.01, respectively. All raw data, statistical representations of the data, and t-test outputs are available in Supplementary Files 1 - 3.

## 3. Results and Discussion

### a. Electron microscopy and EDX

Micrographs were recorded for each biochar at 100x and 1000x magnification. The plant residue biochars RGT and RT exhibited a more heterogeneous composition, exemplified by a broader range of particle sizes, in comparison with the walnut and thermomechanical pulp BC (Figure 1). The proprietary Cool Terra BC featured a more consistent structure, possibly due to post-pyrolysis modification. The micrographs represent a very small sample from each biochar and while more detailed analyses are needed, these results have allowed for the development of several hypotheses regarding how the BC molecular structure might impact various parameters when added to soil. Suffice to say that each feedstock generates biochar with unique microscopic structure, which more than likely imparts different physical and chemical properties (Figure 1).

**Figure 1:**
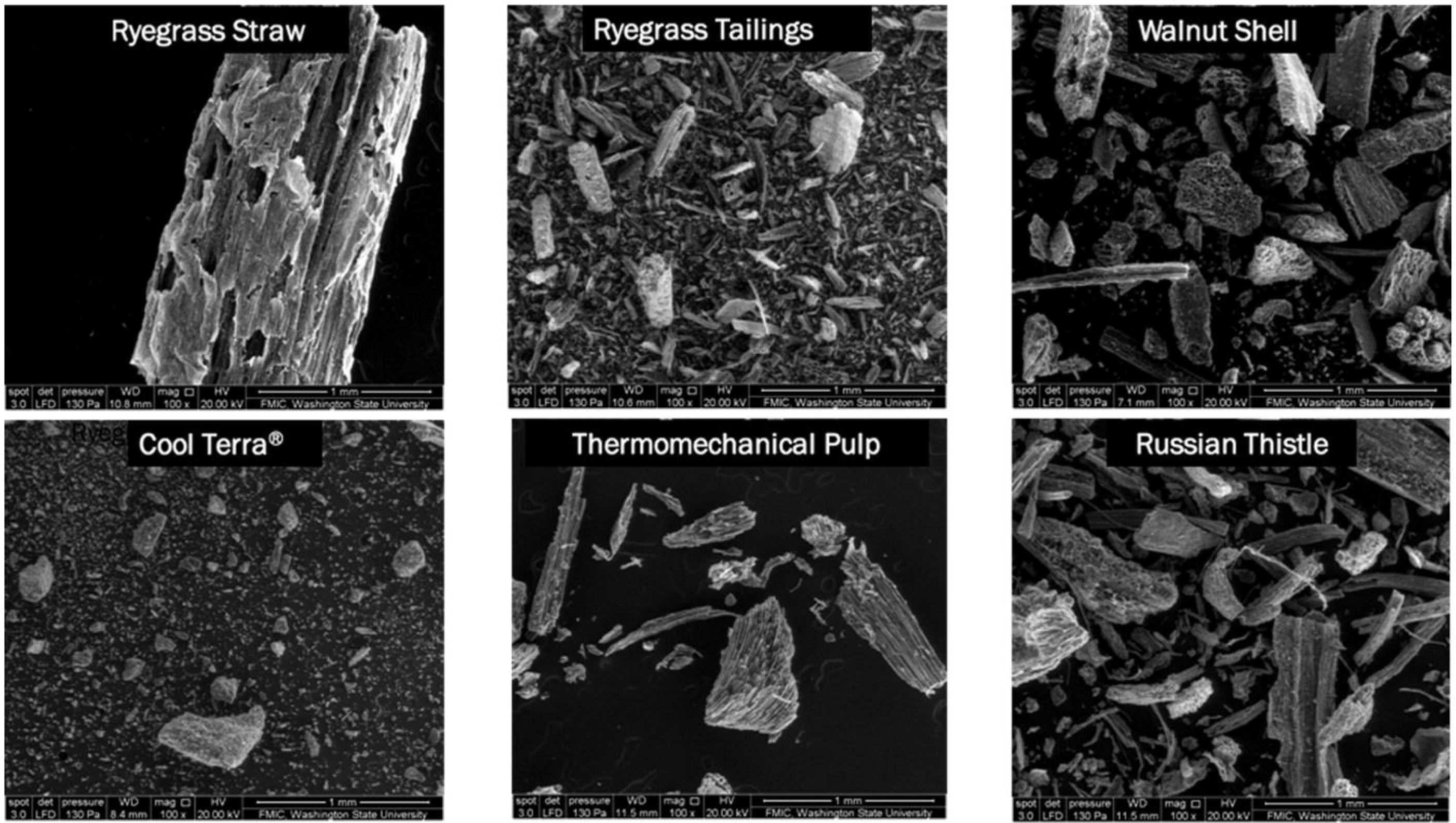
Scanning electron micrographs of different biochars used in this study. (100x magnification and a 1mm scale bar standard). The high heterogeneity of BC is apparent when BC is derived from different feedstocks.

Characterization of all BCs with EDX spectra facilitated qualitative estimation of the specific elements present in each BC. The EDX method is an analytical technique that relies on X-ray excitation and its interaction with a given sample. The unique atomic structure of each element in a sample corresponds to distinctive peaks on the electromagnetic emission spectrum, allowing for chemical and elemental characterization [51]. Nitrogen (N), phosphorus (P), potassium (K), calcium (Ca), and silicon (Si) were the most abundant elements in all BC varieties (Table 1). Sulfur (S) and aluminum (Al) were found in three BCs (RT, Russian thistle, W, walnut, and CT, Cool Terra®), while chlorine (Cl), molybdenum (Mo), magnesium (Mg) and sodium (Na) were only scarcely distributed among the BCs.

Ryegrass tailings (RT)-derived BC contained all analyzed elements except Cl and Na, while the only elements identified in walnut BC were N, P, Ca, and Al (Table 2). While this study used EDX to qualitatively assess BC elemental composition, it is feasible to use this methodology for quantitative elemental analysis [52]. The elemental composition observed is consistent with results of other studies that examined the chemical properties of BCs. These results indicate that feedstocks influence the chemical composition of their biochar derivatives, which vary further based on pyrolysis temperature, and retention time [53].

**Table 2.**
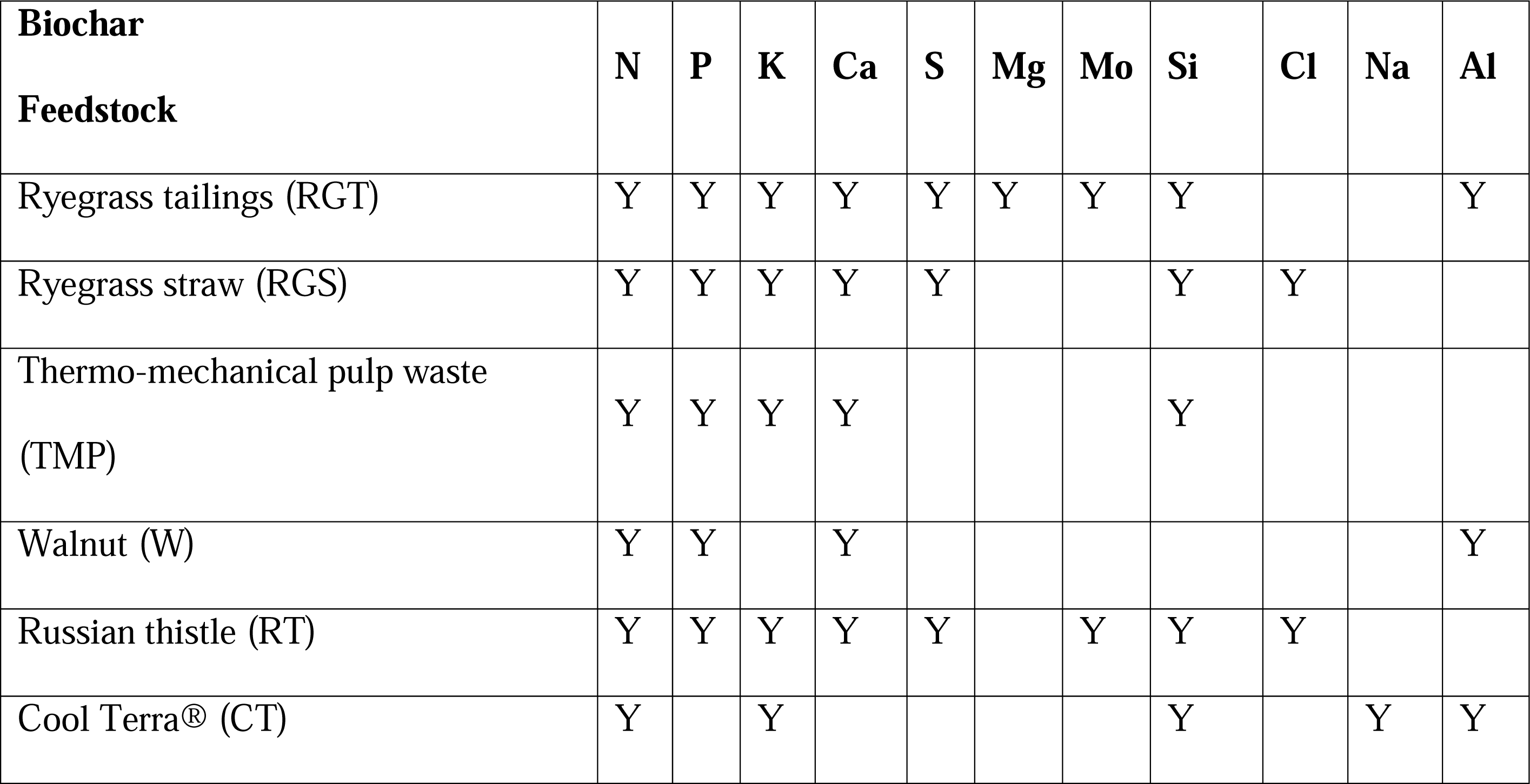
Qualitative elemental composition of different biochars using EDX spectral analysis. Boxes with Y indicate the presence of elements, while blank boxes denote that the element was either not detected or below the detection threshold.

### b. Plant growth parameters

The impact of different biochars on three cultivars of tomato was assessed by quantifying growth and fruit development parameters of the plants grown in the greenhouse, including: shoot dry weight, total fruit weight, and yield per plant.

### i. Shoot Dry Weight

In ‘Oregon Spring,’ decreased shoot dry weight was observed under CT 0.5% (Exp1), RGS 1% (Exp2), and RGT 0.5% (Exp2) BC treatments. However, RGS 1% (Exp1), RGT 1% (Exp1), and TMP 0.5% (Exp1) BC treatments resulted in increased shoot dry weight over control plants (Figure 2A). In ‘Heinz’, increased dry shoot biomass accumulation was observed following all BC treatments except RGS treatments and RGT 1% application in Exp 1. In Exp 2, biochar treatments RGT 0.5%, TMP 0.5% and W 0.5% resulted in decreased shoot dry weight (Figure 3A). For ‘Cobra’, an increase in shoot dry weight was observed in Exp1 with CT, RGS, RGT, and TMP BC applications of 1%, and with 0.5% Russian thistle BC. In Exp2, a decreased in shoot dry weight was recorded with RGS 0.5%, RGT 1%, and TMP 0.5% (Figure 4A).

**Figure 2:**
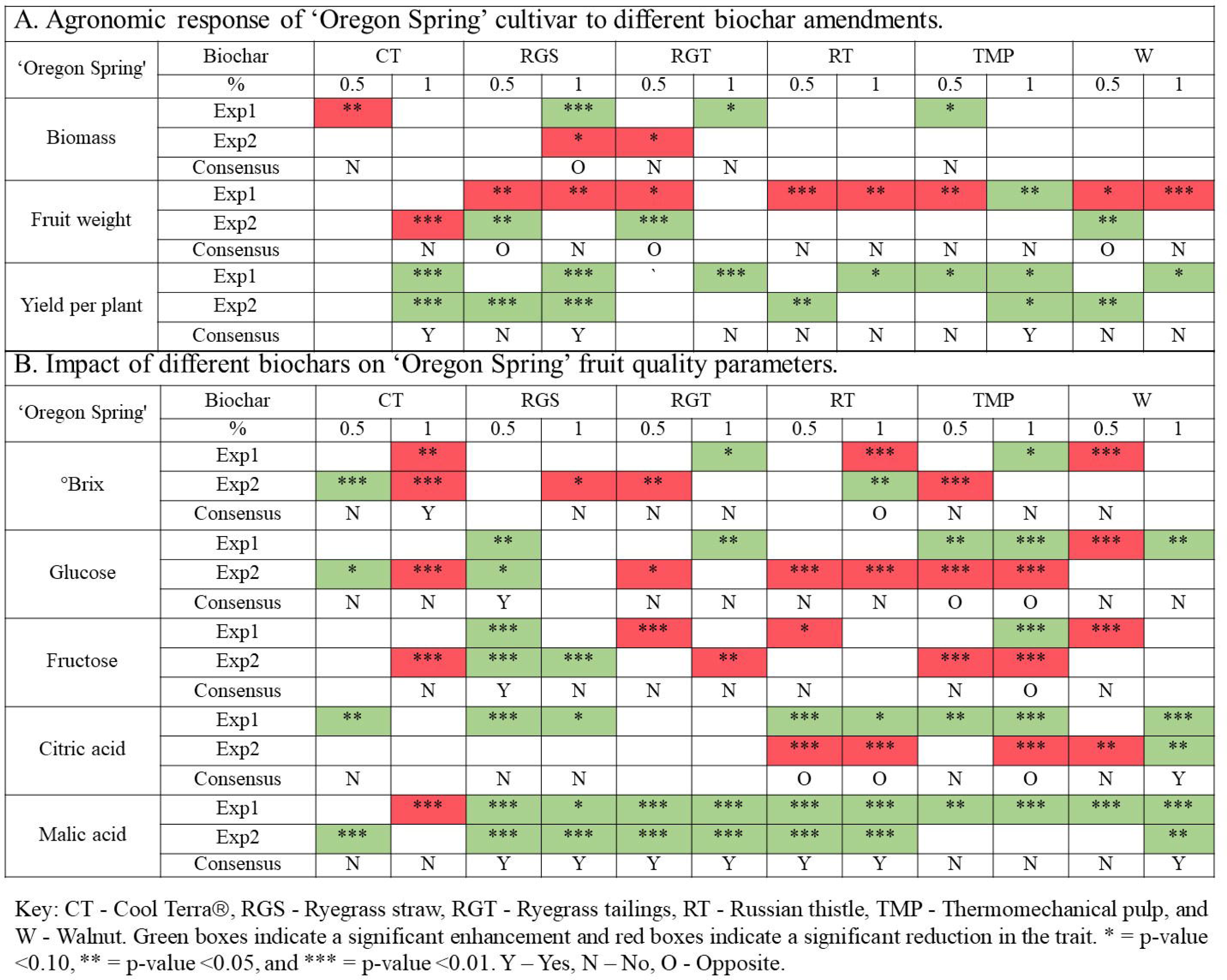
Impact of different Biochars on *Solanum lycopersicum* var. ‘Oregon Spring’ production. A. Agronomic response of ‘Oregon Spring’ cultivar to different BC amendments recorded in terms of biomass, average total fruit weight, and yield per plant (YPP). B. Impact of different Biochars on fruit quality parameters - °Brix, Glucose, Fructose, Citric Acid and Malic Acid.

**Figure 3:**
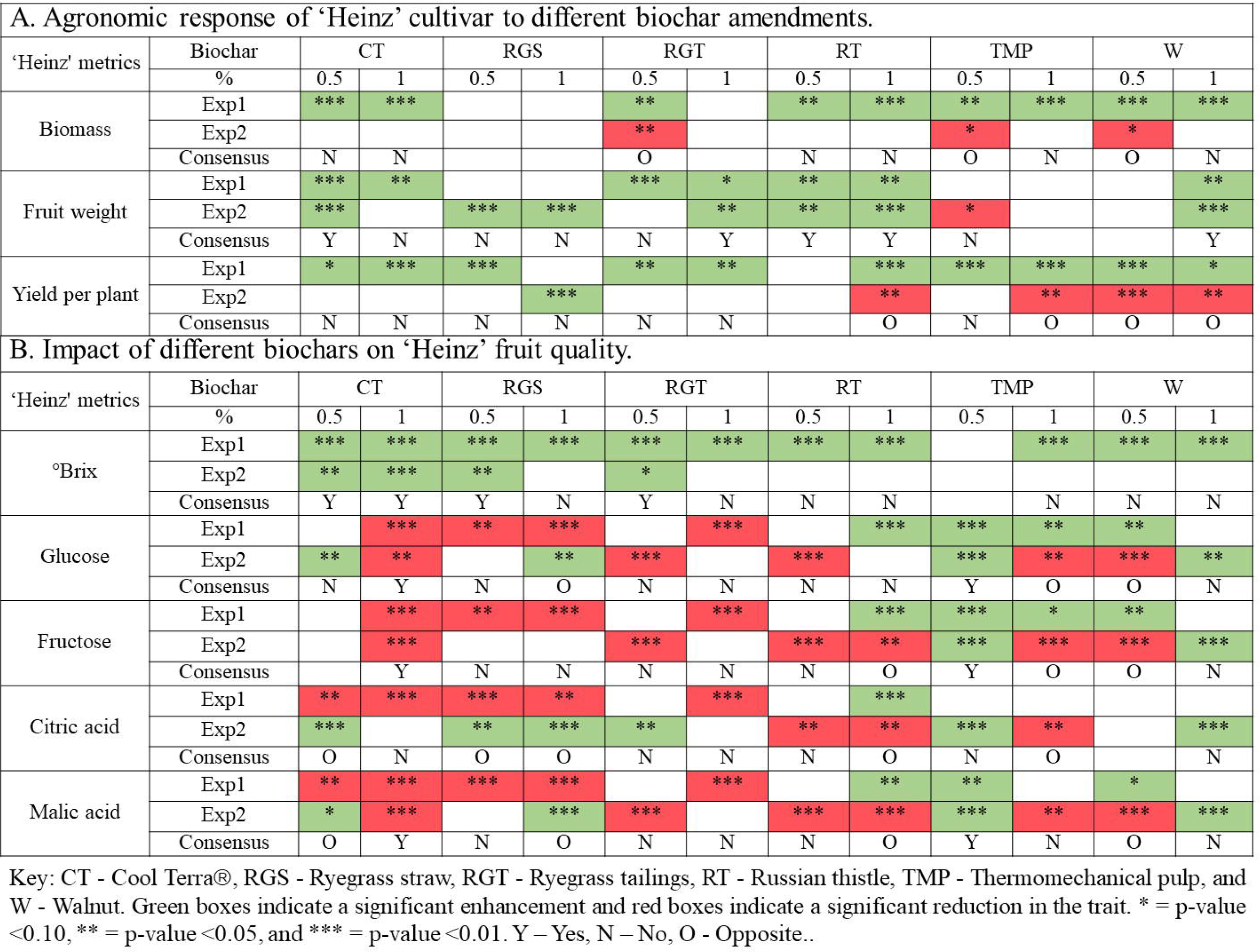
Impact of different Biochars on *Solanum lycopersicum* var. ‘Heinz’ production. A. Agronomic response of ‘Heinz’ cultivar to different BC amendments recorded in terms of biomass, average total fruit weight, and yield per plant (YPP). B. Impact of different Biochars on fruit quality parameters - °Brix, Glucose, Fructose, Citric Acid and Malic Acid.

**Figure 5.**
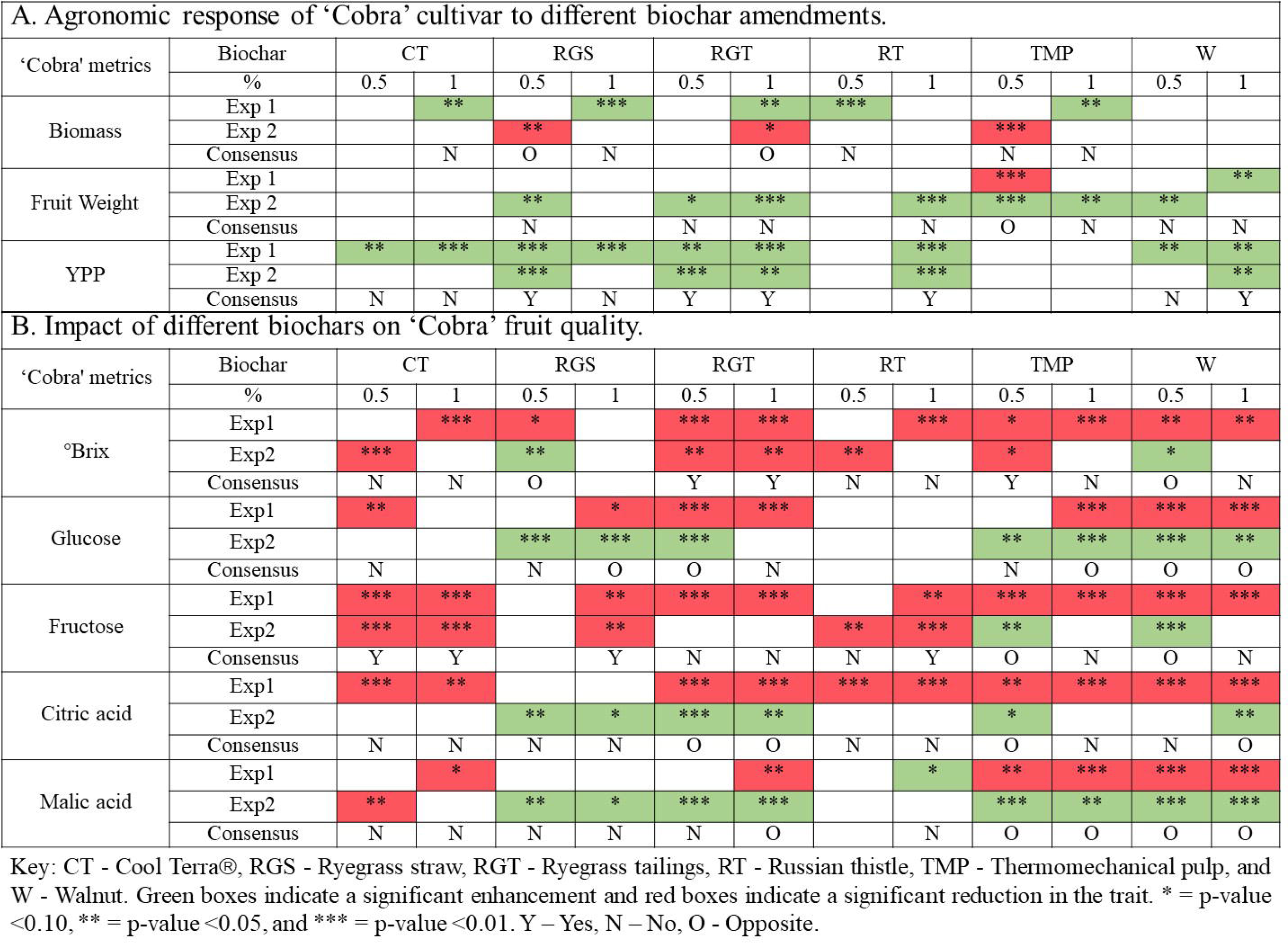
Impact of different Biochars on *Solanum lycopersicum* var. ‘Cobra’ production. A. Agronomic response of ‘Cobra’ cultivar to different BC amendments recorded in terms of biomass, average fruit weight, and yield per plant (YPP). B. Impact of different Biochars on fruit quality parameters - °Brix, Glucose, Fructose, Citric Acid and Malic Acid.

The biomass data across treatments varied between the individual experiments conducted during different times of the year. This was most likely due to the well-documented changes in solar radiation during the year in greenhouse production, and the resulting influence on plant’s photosynthetic performance [54].

In several previous studies, an increase in dry weight was reported following BC application. When wheat bran-derived BC was applied at 14 t/ha rate in tomato production in the field, dry shoot and root vegetative biomass increased, reportedly due to increased soil fertility [32]. Similar results were reported with low-temperature cotton stalk BC in a greenhouse study [55]. A significant increase of up to 52% in shoot dry weight (and 36% increase in root dry weight) was observed in BC-treated plants, in comparison to control plants irrigated with ground water, heavy metal-contaminated water, and sewage water irrigation.

### ii. Fruit weight

#### a. ‘Oregon Spring’

A range of responses was recorded in both experiments under certain treatments with ‘Oregon Spring’ (Figure 3A). Interestingly, the two applications of TMP biochar induced contrasting plant responses in Exp1. The lower dose of TMP 0.5% resulted in fruit with an average mass of 89.9 grams per plant (se +/-4.3) (p<0.05) compared to 99.6 grams (se +/-3.9) in control plants, a 9.7% decrease. The opposite result was obtained with the application of 1% TMP, with the fruit weight increasing 10.5% over control plants to 110.1 grams (se +/-6.6) (p<0.05).

#### b. ‘Heinz’

Fruit weight was significantly increased in multiple biochar treatments for the ‘Heinz’ cultivar (Figure 3A). Average weight of control fruit samples ranged from 42.8 grams (se +/-1) in Exp1 to 49.4 grams (se +/-1.6) in Exp2. The increase in average fruit weight with applications of CT 0.5% ranged from 47.9 grams (se +/-1.7) in Exp1 to 57.5 grams (se +/-2.4) in Exp2 with a p-value of <0.01 for both experiments. This resulted in an increased average fruit weight of 11% in Exp1 and 16% in Exp2. Compared to control samples, RGT 1% treatment also significantly increased average fruit weight in ‘Heinz’ by 4% (p<0.10) in Exp1 and 9% (p<0.05) in Exp2. Additionally, ‘Heinz’ fruit weight increased with both treatments of RT biochar with RT 1% in Exp2 increasing fruit weight by 14% resulting in an average fruit weight of 56.7 grams (se +/-2.1) with significance at p value of <0.01. Walnut biochar at 1% application rate also significantly increased fruit weight in Exp1 to 46.2 grams (se +/-1.8) (p<0.05) and to 57.5 grams (se +/-3.6) in Exp2 (p<0.01), an increase of 7.9% and 16.7%, respectively.

#### c. ‘Cobra’

‘Cobra’ cultivar demonstrated only two significant changes in Exp1 for fruit weight (Figure 4A). A 16% decrease in fruit weight was found following applications of TMP 0.5% (p<0.01), while an 11% increase was shown with W 1% application (p<0.01). In Exp2, both applications of RGT (Exp1, p<0.1; Exp2, p<0.01) and TMP (Exp1, p<0.01; Exp2, p<0.05) resulted in increased fruit weights over control plants in addition to the lower doses of RGS (p<0.05) and W at 0.5% (p<0.05).

Previous studies with *Solanum lycopersicum* cv.’ Brickyard’ in potted bags also showed a different trend. Tomato yield was reported to remain unchanged with 0.5, 1, 2, 4, or 8% BC applications versus controls in trials with ‘Cobra’ cultivar [56,57]. A field study with wheat bran BC applied at 14 t ha–1 reported no impact on fruit weight [32]. The lack of change in average fruit weight most likely indicates that these studies did not have the appropriate biochar type for the specific cultivar to produce any significant effect. In this study, different treatments led to increased average fruit weights observed in ‘Heinz’, ‘Oregon Spring’ and ‘Cobra’, lending support to the original hypotheses.

### iii. Yield Per Plant

In order to evaluate the BC effect on overall crop productivity, average yield per plant (YPP) was recorded. Two of the three cultivars demonstrated an increase in YPP, and no detrimental impact on yield was observed with any biochar treatment.

#### a. ‘Oregon Spring’

Biochar treatments resulted in a significant yield change in ‘Oregon Spring’ cultivar. With applications of CT at 1%, significant increases were found in both experiments, with Exp 2 data indicating a 17% increase (p<0.01) in yield. The average yield was 834.4 grams (se +/-43.9) compared to control plants at 710.3 grams (se +/-33.4). In Exp 1, control plants averaged 634.9 grams (se +/-27.5) and the CT 1% treatment resulted in a significant increase of 12% (p<0.01). Addition of RGS 1% and TMP 1% also significantly increased yields in comparison to the controls in both experiments, especially in Exp1, resulting in a 22% and 12% increase (p<0.01), respectively. An increase in yield was also observed with RT and W biochars in both experiments, with no yield penalties recorded for any BC in either experiment.

#### b. ‘Heinz’

In case of ‘Heinz,’ the YPP data varied across all BC treatments. Contrasting results for both experiments were obtained with 1% concentrations of RT, TMP, and W biochars. In Exp 2, addition of W BC at 0.5% resulted in decreased average yield per plant by 21% (p<0.01). In some cases, increased YPP was observed in Exp 1. Addition of RT 1% and W 0.5% increased yields by 28% and 25%, respectively. This translated to 686.6 grams and 672.2 grams of fruit per plant compared to 535.4 grams in control plants although, the results were reversed in Exp 2.

#### c. Cobra

The YPP in ‘Cobra’ varied across the BC treatments. In Exp 1, control plants averaged 1488.2 grams (se +/-54.9) of fruit while applications of RGT at 1% significantly increased the yield per plant by 20% (p<0.01) to an average of 1788.3 grams (se +/-29.7). While control plants averaged 1053.7 grams (se +/-74.1) in Exp 2, application of RGS at 0.5% resulted in an increase to 1380.7 grams (se +/-126.5), a significant increase of 31% (p<0.01). Additional significant changes were found in Exp 2 for RT 1% applications, for which a yield increase of 25% (p<0.01) was observed. W 1% also raised yield by 19% (p<0.05). TMP-derived biochar at both concentrations had no effect on YPP.

The results of this study demonstrate that different BCs had a positive impact on the productivity of two cultivars, ‘Cobra’ and ‘Oregon Spring’, and only fruit quality metrics in ‘Heinz’ (Figures 2A, 3A, and 4A). Also, a single biochar produced different responses in different cultivars. RGT biochar consistently enhanced yield in ‘Cobra’ but had no effect on the other cultivars. To the best of our knowledge this study is the first to demonstrate an interaction between different feedstock-derived biochars and the genetic background of the plant species.

The observations in this study are in concordance with a recent meta-analysis that summarized the results of 371 independent reports on the effect of biochar on plant productivity and nutrient cycling [44]. While the study found overall positive impact on aboveground biomass (n=67, *P* < 0.01), there were some instances where the BC had a neutral or even a negative impact on plant productivity. In this study, some BC-cultivar combinations enhanced the measured trait, while the majority were neutral, and some produced a negative effect. Interestingly, the experiments yielded contradictory results across the two experiments conducted at different times of the year. Additionally, a recent review on biochar and the effects on agriculture also supports the role of BC as a viable soil amendment to help improve crop productivity while stimulating other soil properties and microbial communities [43].

Previous reports on tomato and other crops have demonstrated mixed results. It was noted that there was an increase in tomato fruit diameter, and a significant yield increase in grape, with BC and compost-amended soils [58]. Additional studies on tomato growth and development with biochar reported similar trends. A field trial with cultivar ‘Trust’ with 10 or 20% (v/v) hardwood BC generated from balsam fir and spruce showed no difference in crop yield [59]. Tomato cultivar cv. 1402 grown in fertigated soilless media also reported no yield increases but did increase plant stature and leaf size. Pepper plants (*Capsicum annuum* L.) reported beneficial yield gains with addition of citrus wood biochar [60]. Enhanced abundance of rhizosphere microbes in addition to a hormesis effect that stimulated plant growth was reported [60].

Results from the above mentioned studies imply that there is a need to further characterize BC-plant interactions. Both the impact of the biological, physical and chemical changes in soil characteristics, and the role of genetic background of the plant will need to be considered if BC is to be deployed widely in agricultural production [43]. The observations summarized in Figures 2, 3 and 4 indicate that different BC treatments generated a unique response in each cultivar and that each cultivar responded uniquely to each BC treatment supporting both hypotheses of this study.

c. Assessment of fruit quality

### i. °Brix

Total Soluble Solids (TSS) assays were conducted for each cultivar. The data indicated mixed results; two of the three cultivars displayed decreased °Brix, with only ‘Heinz’ demonstrating an increased TSS level. The majority of BC treatments had no or negative effect on °Brix.

#### a. ‘Oregon Spring’

A range of TSS response in ‘Oregon Spring’ were observed. Amendment with Cool Terra→ (CT) biochar at 1% resulted in fruit with significantly lower °Brix. In Exp 1, control fruit Brix averaged 5.34 (se +/-0.10) compared to the significantly lowered CT treatment °Brix of 5.04 (se +/-0.14), a 5% decrease (p<0.05). A similar response in Exp 2 resulted in an 8% decrease of °Brix levels in CT 1% fruit versus control fruit. Surprisingly, the lower dose application of CT at 0.5% resulted in the highest TSS level measured in Exp 2, a 8.9% increase, at 6.58 (se +/-0.15) with a p-value of <0.01. These contrasting results necessitate further research to reveal the underlaying plant-BC interaction mechanisms.

#### b. ‘Heinz’

Three BC feedstocks in ‘Heinz’ significantly impacted fruit TSS trends with no detrimental effects measured with any BC treatment or application rate. The data revealed consistently increased °Brix compared to controls with CT 0.5% and 1% (p<0.01), significantly increasing TSS by ∼13% in Exp 1. This resulted in increased °Brix levels of 5.85 (se +/-0.18) in CT 0.5% and 5.84 (se +/-0.17) in CT 1% treatments. More changes were found with RGS 0.5% and RGT 0.5% resulting in °Brix levels increased 10% and 11%, respectively, over controls in Exp 1.

#### c. ‘Cobra’

The ‘Cobra’ cultivar responded comparably with only one biochar type, Ryegrass tailings. The RGT amendment resulted in decreased °Brix at both rates and experimental trials. The largest change was measured in Exp 1 with control fruit °Brix averaging 5.13 (se +/-0.08), while TMP 1% treatment reduced °Brix by 8% to 4.71 (se +/-0.14). Further decreases of TSS were found with all tested biochars in at least one experiment and one concentration.

Only the processing tomato, ‘Heinz,’ revealed increased °Brix concentrations in all biochar treatments (except TMP 0.5%) and in both experiments. Conversely, ‘Cobra’ fruit displayed significantly decreased °Brix levels for four of the six biochars. However, a spectrum of effects was observed for the ‘Oregon Spring’ cultivar. These data supported both of the tested hypotheses.

The beneficial results with ‘Heinz’ indicate this cultivar may be a potential candidate for targeted fruit quality improvement with BC amendment. However, careful consideration of other biochar-cultivar combinations is necessary, as demonstrated by the significantly lowered °Brix levels in ‘Cobra’ cultivar. In a previous study with tomato cultivar ‘Rio Grande’, no substantial changes to the TSS levels were reported when grown in wheat-straw, poplar tree, or olive-residue BC-amended soils at 10% and 20% [61]. Variable response to different BC treatment in terms of °Brix was observed in this study as well. The multi-variable BC-plant-soil interactions along with the genetic background of the plant most likely influences TSS levels.

### ii. Sugars

Producing flavorful tomatoes could be an advantage to producers, processors, and consumers alike. Sugars were quantified with HPLC to determine the carbohydrate load in tomato fruit in response to BC amendment.

#### a. ‘Oregon Spring’

A wide range of responses were recorded in ‘Oregon Spring’ cultivar in response to various BC treatments (Figure 2B). Significant changes in glucose and fructose levels were recorded with RGS treatment at 0.5% in both experiments. In Exp 1, glucose concentrations were significantly increased by 18% (p<0.01) from 14.35 μg/μL (se +/-0.80) in control plants to 16.98 μg/μL (se +/-1.26) in RGS treated plants. In Exp 2, fructose levels were also increased by 25% in RGS 0.5% treatment as indicated by the increase from 38.58 μg/μL (se +/-1.4) in control fruit to 48.28 μg/μL (se +/-3.37) (p<0.01).

#### b. ‘Heinz’

Various significant differences were observed in ‘Heinz’ fruit carbohydrate levels (Figure 3B). ‘Heinz’ demonstrated significantly increased glucose and fructose levels over control plants, with TMP 0.5% BC treatment (p<0.01). In Exp 1, TMP 0.5% increased fruit glucose levels by 27% from 13.06 μg/μL (se +/-0.95) in control fruit to 16.7 μg/μL (se +/-1.67). Fructose levels responded similarly, with a 30% increase from 25.52 μg/μL (se +/-2.16) in control fruit to 33.28 μg/μL (se +/-2.71) (p<0.01). However, applications of CT at 1% significantly decreased glucose by 24% (p<0.01) and fructose by 27% (p<0.01) in Exp 1, with similar trends in Exp 2.

#### c. ‘Cobra’

Although significant differences were found in glucose levels, no consistent results were observed between the BC treatments in ‘Cobra’ (Figure 4B). Fructose levels were impacted by several biochar feedstocks and demonstrated significant decreases with 1% applications of CT, RGS, and RT. A steep decline in fructose was observed with both applications of W BC in Exp 1 lowering the levels by 25% and 27% compared to control fruit. The CT treatments incrementally lowered fructose levels with increasing BC rates from 30.18 μg/μL (se +/-1.91) in the 0.5% treatment (13.8% decrease) to 27.78 μg/μL (se +/-3.54) in the 1% treatment (20.9% decrease) compared to 35.02 μg/μL (se +/-1.14) in the control fruits (p<0.01).

The impact of BC on carbohydrate levels in the three cultivars was highly diverse. In a previous study demonstrated that tomato TSS was negatively affected by olive-residue BC, indicative of BC-specific effects on tomato quality. Another study showed lower temperature (300 C°) BC treatments resulted in increased sugar levels in ‘Micro-Tom’ tomato in a pot study with BC derived from bamboo feedstock [62]. These results further indicate a BC specific effect on fruit quality that is also dependent on the genetic background of the cultivar.

### iii. Organic acids

Similar to other traits, a range of responses was recorded for the quantified organic acids. ‘Oregon Spring’ demonstrated a strong response to BC treatments, especially in terms of malic acid (MA) levels, which increased in both the experiments and rates using RGS, RGT, and RT BC. Compared to controls, plants amended with RGT 1% resulted in significantly higher MA production as shown in Exp 1 (63%) and Exp 2 (30%). Additionally, in Exp 1, RT 0.5% amendment resulted in a significant increase of 80% in MA levels from 0.89 μg/μL (se +/-0.07) in control fruits to 1.60 μg/μL (se +/-0.15) in treated fruits (p<0.01). Other BC treatments in Exp 1 also significantly (p<0.01) altered MA levels in ‘Oregon Spring’ as W 1% and TMP 1% increased MA by 74% and 80% while CT 1% decreased MA by 18% to 0.73 μg/μL (se +/-0.05) (Figure 2B).

A similar pattern of variable but significant changes was recorded in the ‘Heinz’ cultivar. In Exp 1, applications of RT BC at 1% increased citric acid levels by 66% (p<0.01) and malic acid concentrations by 58% (p<0.05). A significant difference (p<0.01) was found with CT 1% treatment, which reduced the MA levels from 1.08 μg/μL (se +/-0.09) in control fruit to 0.63 μg/μL (se +/-0.10), a 41% decrease in Exp 1. Conversely, MA levels in Exp 2 significantly increased (p<0.01) by 25% with TMP 0.5% treatment compared to control fruit (Figure 3B).

In the ‘Cobra’ cultivar citric acid (CA) and malic acid (MA) levels decreased significantly (p<0.01) in Exp 1 with 0.5% Walnut biochar: CA decreased 27% and MA decreased by 16%. No decreases were observed in Exp 2 with W BC but RGT BC at 0.5% increased CA (21%) and MA (11.5%) concentrations significantly (p<0.01) (Figure 4B).

A previous study showed that tomato fruit quality, specifically CA, did not statistically improve between BC treatments and even showed a significant decrease with 10% olive-residue BC [61]. The generally positive response to BC amendment in the ‘Oregon Spring’ cultivar in terms of organic acids compared to the other cultivars most likely indicates a more favorable plant-soil-genetic background interaction. These data support both hypotheses as each biochar affected fruit quality differently, and each cultivar had a unique response to each BC (Figure 2B, 3B, and 4B).

## 4. Conclusion

The data presented in this study supports both the hypotheses, 1.) Biochars derived from different feedstock sources will produce unique phenotypes in a single cultivar of tomato, and 2.) a single feedstock-derived biochar will produce different phenotypes in each of the three tomato cultivars. The results indicate towards future experiments to focus on understanding all BC-related variables, the significance of their contribution individually and in an interactive context when added to soil. There is a need to adopt a customized approach for BC application in order to enhance yield and quality of the crop [24,33,43,63]. Future BC studies should evaluate multiple crop cultivars in conjunction with different classes of biochar (ex. manure, hardwood, or crop residue), to dissect the nature of the complex interactions.

In summary, in ‘Oregon Spring’, a preferred tomato variety for backyard production, the yield per plant and malic acid were seen to be enhanced, and there was a general consensus between the two experiments for these two traits. Overall, CT 0.5%, and RGS 0.5% treatments were most suitable for enhancing fruit quality traits. The results from the processing tomato, ‘Heinz,’ were different. Most BC-treatments enhanced growth and development traits. However, °Brix and other fruit quality traits were negatively impacted except for TMP 0.5% treatment. ‘Cobra’, a variety bred for greenhouse production, showed enhanced yields in all experiments and most BC combinations; however, fruit quality traits varied across all BC treatments. While additional experimentation is required to understand the wide-ranging variability in responses, several possible variables can influence the outcomes, including: feedstock, potting mix, biochar characteristics, microcosm, environmental factors, production management and the genetic background of the plant. The observations recorded in this study should be considered with the caveat that the experiments were conducted in a greenhouse. Use of potting mix eliminated all the soil-related dynamics that may have influenced the agronomic performance and fruit traits. Nevertheless, the study demonstrates that the genetic background of the plant is an important variable. Prospective field evaluation of biochar should include different cultivars of the species being tested. Moreover, the productivity of future agroecosystems will be measured by the intensity of current attempts to improve soil health; therefore, methods that improve and maintain soil health should be incorporated and evaluated rigorously.

## Declarations

### Ethics approval and consent to participate

Not applicable

### Consent for publication

Not applicable

### Availability of data and material

The datasets supporting the conclusions of this article are included within the article and its additional files.

### Author Contributions

Conceptualization, AD, DI; Methodology, DI, ET, RG, AD; Formal Analysis, ET, RG, AD; Resources, AD; Writing – Original Draft Preparation, ET, AD; Writing – Review & Editing, ET, RG, AD; Supervision, AD; Funding Acquisition, AD,

## Acknowledgments

The authors thank Dr. Seanna Hewitt and Evan Stowe for their critical reading of the manuscript and their feedback. The authors also thank Milica Radanovic for help with statistical analysis. The authors are thankful to Dr. Seanna Hewitt and Robert Nica for designing the graphical abstract.

## Abbreviations

BC: Biochar
SOM: soil organic matter
SOC: soil organic carbon
CT: Cool Terra®
RGS: Ryegrass straw
RGT: Ryegrass tailings
RT: Russian thistle
TMP: Thermomechanical pulp

W: Walnut
SEM: Scanning Electron Microscopy
EX: energy-dispersive X-ray spectroscopy – EDX
w/w: weight/weight
High Pressure Sodium: HPS
total soluble solids: TSS
N: nitrogen
P: phosphorus
K: potassium
Ca: calcium
S: sulfur
Mg: magnesium
Mo: molybdenum
Si: silicon
Cl: chlorine
Na: sodium
Al: aluminum

